# A precise IDMS-based method for absolute quantification of phytohemagglutinin, a major antinutritional component in common bean

**DOI:** 10.1101/2023.12.07.570538

**Authors:** Lan Li, Zhanying Chu, Kang Ning, Manman Zhu, Rui Zhai, Pei Xu

## Abstract

Phytohemagglutinin (PHA), a natural tetramer comprising PHA-E and PHA-L subunits that preferentially bind to red and white blood cells, respectively, constitutes a significant antinutritional and allergenic factor in common bean seeds. The accurate measurement of PHA content is a prerequisite for ensuring food safety inspections and facilitating genetic improvements in common bean cultivars with reduced PHA levels. Currently, mainstream methods for PHA quantification involve hemagglutination assays and immunodetection, but these methods often require fresh animal blood and lack specificity and accuracy. In this study, we present a novel LC-MS/MS-based method for PHA quantification, leveraging the advantages of isotope dilution mass spectrometry (IDMS). Two signature peptides each for PHA-E and PHA-L, along with a common signature peptide, were identified and employed for quantification, allowing differentiation between PHA-E and PHA-L subunits. The incorporation of amino acid analysis-isotope dilution mass spectrometry (AAA-IDMS) enabled precise determination of the synthetic signature peptides’ purity during measurement, enhancing metrological accuracy. In addition, the TCA-acetone protocol was established as the optimized method for total protein extraction from dry bean seeds. Quantitative analysis of PHA-E and PHA-L subunits in six common bean varieties using the developed method demonstrated excellent linearity (r > 0.999), sensitivity (limit of detection and quantitation as low as 2.32 ng/mg and 7.73 ng/mg, respectively), recovery (94.18-104.47%), and repeatability (relative standard deviation < 3.45%). This method has the potential to serve as a standard for measuring PHA contents in common beans and other agricultural products containing PHA.

## 1 Introduction

Common bean (*Phaseolus vulgaris* L., 2n=22), also known as French bean or kidney bean, belongs to the *Phaseolus* genus in the *Leguminosae* family. With a genome size of approximately 600 Mb, common bean is among the most widely cultivated food legume worldwide (Bitocchi et al., 2017). This crop is a rich source of protein in both its pods and seeds, providing up to 70% of the total protein requirement for local residents in Latin America and Africa, where affordable animal proteins are scarce (Petry et al., 2015). Furthermore, it is an excellent source of essential micronutrients such as iron, phosphorus and magnesium (Celmeli et al., 2018). However, the sustainable development of the common bean industry is facing challenges due to concerns about the safety of consuming its seeds. Intake of inadequately processed bean seeds or pods with seeds has been associated with allergic reactions and digestive system symptoms, such as nausea, vomiting and dizziness (Muramoto, 2017). Consequently, common bean has been included in the list of food safety priorities in many countries (Ogawa & Date, 2014; Rodhouse et al., 1990). Research has unveiled that phytohemagglutinin (PHA), a seed storage protein accounting for up to over 10% of the total protein content in seeds, is the primary cause of adverse reactions linked to common beans (Campos-Vega et al., 2010).

PHA belongs to the lectin protein superfamily, which is ubiquitous across bacteria, fungi, plants and animals (Tsaneva & Van Damme, 2020). PHA generally consists of two polypeptide subunits known as PHA-L and PHA-E, which exhibits preferential binding characteristics to red blood cells and white blood cells, respectively, attributed to the presence of specific glycans chains attached to the polypeptides’ surfaces. The PHA-L subunit has a relative molecular weight of about 30 kDa, while PHA-E subunit weights approximately 34 kDa (Leavitt et al., 1977; Einas Hamed et al., 2017). The natural PHAs are tetramers randomly composed of five isomeric forms, including E4, E3L1, E2L2, E1L3, and L4 (He et al., 2018). In addition to agglutinating blood cells, PHA can damage intestinal epithelial cells and bind with digestive enzymes, causing a series of gastrointestinal discomfort symptoms (Miyake et al., 2007). It has also been observed to bind with the immunoglobulin IgE, confirming its allergenicity (Barre et al., 2020).

There are currently a few methods available for determining the content of PHA, including hemagglutination and enzyme-linked immunosorbent assay (ELISA). However, hemagglutination assay requires fresh animal blood and its detection sensitivity and accuracy are low. In addition, because of the existence of glycan chains on the surfaces of many types of lectins other than PHA, the hemagglutination assay lacks specificity (Zhao et al., 2019). ELISA, based on the specific binding of antigen and antibody, is a widely used method for allergen quantification. Nevertheless, it is also prone to cross-reactivity (Osborn et al., 1988) and currently there is a scarcity of commercially available ELISA kit dedicated to PHA detection. Additionally, variations in antigen and antibody production batches necessitate calibration before use, which compromises the reliability of comparing results across different experiments or laboratories (Aydin, 2015).

Liquid chromatography-mass spectrometry (LC-MS) is a powerful tool that is widely used for the identification and quantification of proteins in biological samples. In the case of proteins with large molecular weights, trypsin can be employed to enzymatically digest the protein into smaller peptides with relatively lower molecular weights. From these peptides, peptide segments unique to the protein, known as signature or characteristic peptides, can be selected for quantitative determination of the protein content. The LC-MS-based protein quantification method offers high sensitivity, specificity, accuracy, and reproducibility compared to other quantitative techniques. Moreover, by introducing known amounts of isotopically labelled signature peptides as internal standards in the samples, the foundation for the IDMS method is established, wherein the mixing of the isotopic labeled peptide with the sample effectively eliminate errors of system and improve the accuracy of quantification. Unlike conventional analytical methods that rely on signal intensity, this approach employs signal ratios. This combined quantification strategy, known as isotope dilution mass spectrometry (IDMS), assists in mitigating errors arising from random and systematic variations, and is regarded among chemistry measurement methods of the highest metrological standing (Milton & Wielgosz, 2000).

He et al. (2015) employed LC-MS/MS analysis following in-gel tryptic digest to identify a 31 KDa protein isolated from the black turtle bean as a PHA-E protein. However, by far, LC-MS-based quantification method for PHA has not been established. In this study, we developed a novel LC-MS/MS-based method for the identification and absolute quantification of PHA. This method can not only distinguish the PHA subunits but has a compelling precision and repeatability in quantification.

The method is poised to facilitate food safety inspection and enhance genetic improvement for common bean cultivars with depleted PHA levels.

## 2 Materials and Methods

### 2.1 Reagents

The following chemical reagents were utilized in the study: Formic acid (LC-MS grade), trifluoroacetic acid (TFA, LC-MS grade), acetonitrile (LC-MS grade) and acetone (HPLC grade) were purchased from Thermo Fisher Scientific (USA). Trypsin, dithiothreitol (DTT), iodoacetamide (IAA), ethylene diamine tetraacetic acid (EDTA), sodium dodecyl sulfate (SDS), phenyl methane sulfonyl fluoride (PMSF), trichloroacetic acid (TCA), ammonium bicarbonate (LC-MS grade) were purchased from Sigma-Aldrich (USA). Tris-HCl, urea, thiourea, hydrochloric acid were products of Macklin (Changhai, China). Tris-buffered phenol was purchased from Damao Chemical Reagent Co., Ltd (Tianjing, China). Peptides and stable isotope-labeled signature peptides for the PHA-E and PHA-L subnuits (see also section 2.3) with the sequences GNVETNDVLSWSFASK, SVLPEWVSVGFSATTGINK, GGLLGLFNNYK, TTTWDFVK, TSFIVSDTVDLK were sourced from Genscript Biochem. Co, Ltd. (Nanjing, China). PHA standard protein were purchased from Thermo Fisher Scientific (USA). L-valine (Val, GBW09236, 99.4%), L-phenylalanine (Phe, GBW09235, 99.8%), L-leucine (Leu, GBW09237, 99.7%), L-isoleucine (Ile, GBW09238, 99.7%) were obtained from the National Institute of Metrology (Beijing, China). The isotope-labeled aminos (AAs*), including ^13^C_6_-Leucine (Leu*, ≥ 99%), ^13^C_9_-phenylalanine (Phe*, ≥ 99%), ^13^C_5_-valine (Val*, ≥ 99%), ^13^C_5_-isoleucine (Ile*, ≥ 99%) and ^13^C_5_-proline (Pro*, ≥ 99%) were purchased from the US Cambridge Isotope Laboratory (USA).

### 2.2 Analytical instruments

Two different instruments were utilized for analysis. The concentration of amino acids and unique peptides was measured using an ultrahigh-performance liquid chromatograph tandem triple quadrupole mass spectrometer (Vanquish UHPLC-TSQ Altis MS, ThermoFisher, USA). To confirm the peptide structure, a linear quadrupole ion trap mass spectrometer (LTQ Orbitrap Elite MS, ThermoFisher, USA) was employed.

### 2.3 Screening of signature peptides

The protein sequences of PHA-L (Entry ID: P05087) and PHA-E (Entry ID: P05088) were retrieved from UNIPROT (https://www.uniprot.org) for further analysis. These protein sequences were then subjected to simulated digestion with trypsin. From the digested peptides, we selected those with a sequence length between 7 and 20 amino acids, m/z values below 1250, and no cysteine (C) or methionine (M) residues. We further considered criteria such as the absence of internal trypsin cleavage sites and the absence of continuous arginine (R) or lysine (K) residues at the restriction site. The peptides meeting the aforementioned requirements were used as queries to search against the *Phaseolus vulgaris* protein library (ID 3885) in UNIPROT, and those hitting only the PHA-E and/or PHA-L proteins were retained as putative signature peptides.

### 2.4 Amino acid analysis (AAA)-IDMS

To ensure a more accurate quantification, the purity of the synthesized peptides was determined by isotope dilution mass spectrometry based on AAA, and IDMS was analyzed under the MRM mode of ultra-high-performance liquid chromatography in tandem with triple quadrupole mass spectrometer. 6M HCl was used to completely hydrolyze the peptides into amino acids. Following collision energy (CE) and radio frequency voltage (RF Lens) optimization, the amino acids were separated on a ACQUITY UPLC^®^ HSS T3 (1.8 μm*100*2.1 mm) column with a mobile phase of water to acetonitrile to FA at 98:2:0.8 (v:v:v). The injection volume was 10 μL and an isocratic elution flow of 0.25 mL/min was used.

### 2.5 Sample preparation

#### 2.5.1 Sample grinding

Raw seeds from six various genotypes of common bean sourced in China, namely JH-40, JH120, JH266, JH227, JH231, JH301 were ground using a high-throughput tissue grinder (Ningbo Scientz Biotechnology Co., Ltd, China). The resulting powder was subsequently frozen at -80℃ until analysis.

#### 2.5.2 Buffers preparation

Wash buffer I: wash buffer I contains 10 mM Tris-HCl (pH 8.0), 1 mM EDTA and Tris-buffered phenol. This solution was stored at 4℃ for no more than a month.

Trichloroacetic acid (TCA), 10% (wt/vol), in acetone: 1.0 g of TCA was dissolved in 9ml of acetone, and 5mM DTT was added. The solution was kept at − 20℃.

SDS extraction buffer: this buffer contains 1% (wt/vol) SDS, 0.1 M Tris-HCl (pH 8.8), 1 mM EDTA, 2 mM PMSF, and 10mM DTT.

Acetone, 80% (vol/vol), in water: 8ml of acetone was mixed with 2 ml of deionized water, with the addition of 5mM DTT. The solution was kept at − 20℃.

#### 2.5.3 Extraction process

Three classical methods for extracting plant proteins were employed and compared in this study (Wu et al., 2014).

TCA/acetone precipitation involved the following main steps: A) The powder was homogenized with pre-cooled SDS extraction buffer at 4℃. B) Centrifugation at 15,000g for 5 minutes at 4℃ to collect the supernatant. C) Addition of 10% TCA-acetone to the supernatant, followed by swirling and placing on ice for 5 minutes. D) Centrifugation at 15,000g for 5 minutes at 4℃ to collect the precipitate. E) Cleaning the precipitate with pre-cooled pure acetone, followed by centrifugation at 15,000g for 5 minutes at 4℃. F) Cleaning the precipitate with pre-cooled 80% acetone, followed by centrifugation at 15,000g for 5 minutes at 4℃. G) Air-drying the precipitate (Niu et al., 2018).

Phenol extraction involved the extraction of proteins from an aqueous solution using phenol (Alara et al., 2021; Faurobert et al., 2006). The main steps include: A) the powder was homogenized with pre-cooled SDS extraction buffer at 4℃. B) Centrifugation at 15,000g for 5 minutes at 4℃ to collect the supernatant. C) Addition of the same volume of tris-buffered phenol used in the previous step. After being thoroughly mixed and centrifuged at 15,000g for 5 minutes at 4℃, the light-yellow phenol phase was taken. D) Addition of an equal volume of wash buffer I to the phenol phase and well shaking (3-5 minutes). Following centrifugation at 15000g × 5min × 4 ℃, the upper phenol phase was collected. E) Addition of five times the volume of methanol with 0.1M ammonium, well mixing, and leaving at -20 °C for 30 minutes. The extracted proteins in sediments were collected. F) The precipitations were washed with 1.5ml methanol (containing 0.1M ammonium acetate), then centrifugated at 15000 g × 4℃ × 5 min. G) Cleaning the precipitate with 1.5ml of 80% acetone, centrifuging at 15000 g × 4℃ × 5 min, which was repeated twice.

TCA/acetone/phenol method combines the advantages of the above two methods, enabling the removal of non-protein compounds and selective dissolution of proteins (Wang et al., 2003). The main steps include: A) The powder was homogenized with pre-cooled 10% TCA/acetone at 4 ℃. B) Centrifugation at 15,000g for 5 minutes at 4℃ to collect the sediment. C) The precipitate was suspended again in pre-cooled 10% TCA/acetone, followed by centrifugation and collection of the precipitate. This process was repeated until the sediment turned white. D) The precipitate was suspended again in pre-cooled acetone, followed by centrifugation and collection of the sediment. F)

The sediment was air-dried. G) Using SDS extraction buffer homogenate for resuspension, followed by incubation at 60-70℃ for 1-2 hours. H) Centrifugation was performed, and the supernatant collected. I) Tris-buffered phenol was added, and after thorough shaking, centrifugation was performed to collect the phenol phase. J) Precipitation was carried out using methanol plus 0.1M ammonium acetate. K) The protein pellet was washed and air-dried (Wu et al., 2014).

#### 2.5.4 Digestion procedures

The extracted proteins were resuspended with 1.5 mL of 8M urea. The protein concentration was determined using a BCA Protein Assay Kit (Beyotime Biotechnology, China). The protein solution was diluted with 50 mM NH_4_HCO_3_ containing 10 mM DTT, followed by incubation at 95°C for 10 minutes. Alkylation was carried out in the dark for 30 minutes at room temperature using 50 mM iodoacetamide (IAA) in 50 mM NH_4_HCO_3_. DTT was added at room temperature for 10 minutes to neutralize any excess IAA. Finally, trypsin was added at a trypsin-to-protein ratio of 1:5 (w/w), and incubation was conducted at 37°C for 20 hours.

#### 2.5.5 Desalting

Digested samples were adjusted to pH < 2 using 0.1% TFA to terminate the reaction and then desalted with Waters Sep-Pak C18 Cartridges (50 mg), which were activated using 1 mL acetonitrile (ACN) and 1 mL 0.1%TFA. The enzymolysis product of the redissolved protein was transferred to the desalting column, and the process was repeated twice. Desalinate twice with 1 mL 0.1% TFA aqueous solution. Finally, 1mL ACN/0.1% TFA (50/50, v/v) was used to collect peptides.

### 2.6 Liquid chromatography and mass spectrometry analyses

For the select of the signature peptides, the peptide solution was separated by UHPLC with Waters ACQUITY UPLC®HSS T3 (1.8 μm*100*2.1 mm) column, with deionized water (0.1% FA) and acetonitrile (0.1% FA) as the mobile phase A and B, respectively. The LC gradient was set as follows: 15% B from 0 to 5 min, then increased to 40% B within 36 min, 40-100% B from 36 min to 45 min, next returned to 15% B from 45 min to 56 min, finally equilibrated with 15% B for 9 min. The TSQ Altis were used with m/z range of MS acquisition from 300 to 1650.

Multi-response monitoring mode (MRM) in LC-MS/MS was used for accurate quantification of the PHA in common bean. The signature peptide solution was separated by UHPLC with Waters ACQUITY UPLC®HSS T3 (1.8 μm*100*2.1 mm) column. After heat drying, the PHA enzymatic hydrolysis product was dissolved in 200 μl water-0.1% FA and 10 μl of sample was loaded. The content of the mobile phase A and B were water-0.1% FA and ACN-0.1% FA, respectively. The flow rate was 0.2 ml · min^-1^. The elution gradient was: 0 - 3 min, 12% B; 3-6min, 23 % B; 6 - 7 min, 23% B; 7-10min, 27% B; 10-15min, 38% B; 15 - 15.1 min, 90% B; 15.1-18min, 90% B; 18-18.1min, 12% B; 18.1-27min, 12% B. Mass spectrum conditions: MRM mode was selected, ESI source spray voltage was 3.5 kV, and ion transfer tube temperature was 325 °C. The collision energy was 13.75V and the cluster removal voltage was 44V. The mass spectrum data were collected by the Xcaliblur software and analyzed using Thermo TraceFinder.

### 2.7 Estimation of recovery rate

Known amounts of standard proteins PHA-E and PHA-L were added to the sample, and the amount added was denoted as A. After processing according to the sample preparation method 2.5, the total amount of PHA-E and PHA-L protein measured was denoted as B, the sample without standard protein actually measured C. Recovery rate was assessed by formula: recovery rate =(A-B/C)×100%.

### 2.8 Limit of detection (LOD) and limit of quantitation (LOQ)

The performance of the method was evaluated by LOD and LOQ. The limits of detection and quantitation are the analyte concentrations at which the signal-to-noise ratios are ≥ 3 and ≥ 10, respectively.

### 2.9 Assessment of precision

To assess intra-day precision, the identical sample was injected 6 consecutive times within a day to determine the intra-day relative standard deviation (RSD). For assessing inter-day precision, the identical sample was measured every 2 hours for three times in a day, and measurement was performed over three days to calculate inter-day RSD.

## 3. Results and discussion

### 3.1. Identification of signature peptides for PHA quantification

According to the screening principle of section 2.3, we identified a total of 5 signature peptide segments for PHA-E, 6 signature peptide segments for PHA-L, and 2 signature peptide segments as common for both chains (Table S1). We then conducted LC-MS/MS analysis with the peptides to determine the best-performing signature peptides. As a result, 6 peptides were found to exhibit sharp and high response intensity peaks (Fig. 1). These peptides were subsequently synthesized and employed as the quantitative peptides in following steps, which were also validated using LTQ Orbitrap Elite (Fig. S1). The optimized MRM parameters for measuring these signature peptides are listed in Table 1.

**Fig. 1.**
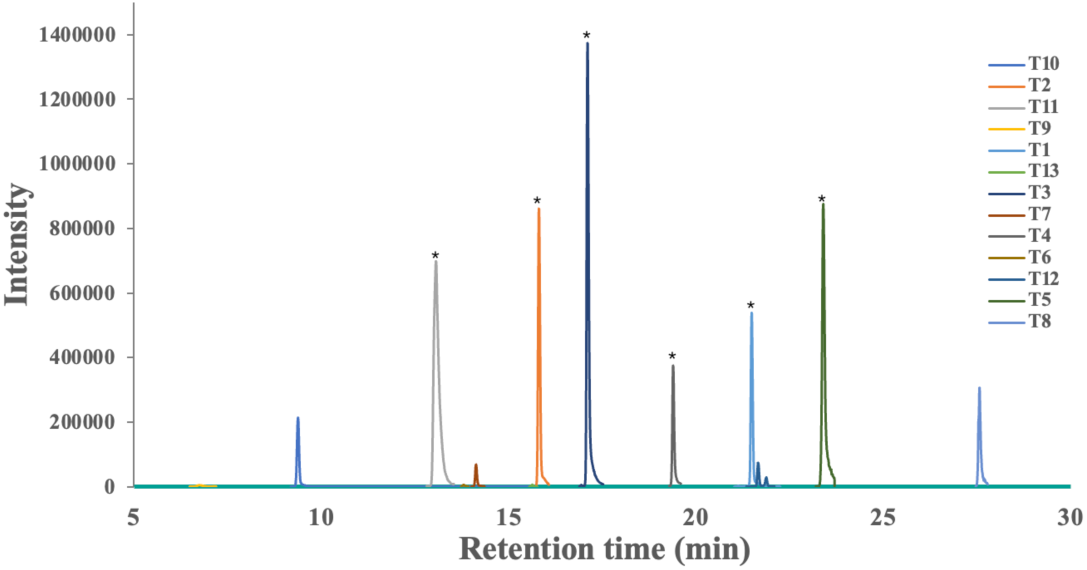
Peptide mass spectra for the 13 candidate signature peptides. T1: GGLLGLFNNYK; T2: TTTWDFVK; T3: TSFIVSDTVDLK; T4: GNVETNDVLSWSFASK; T5: SVLPEWVSVGFSATTGINK; T6: LTNVNDNGEPTLSSLGR; T7: GENAEVLITYDSSTK; T8: SVLPEWVIVGFTATTGITK; T9: LTNLNGNGEPR; T10: DWDPTER; T11: HIGIDVNSIR; T12: LSDGTTSEGLNLANLVLNK; T13: FNETNLILQR. The six peptides showing sharp and high peaks are marked with asterisks. Note that the peptide HIGIDVNSIR (T11) was not successfully synthesized and thus not included in following steps.

**Table 1.**
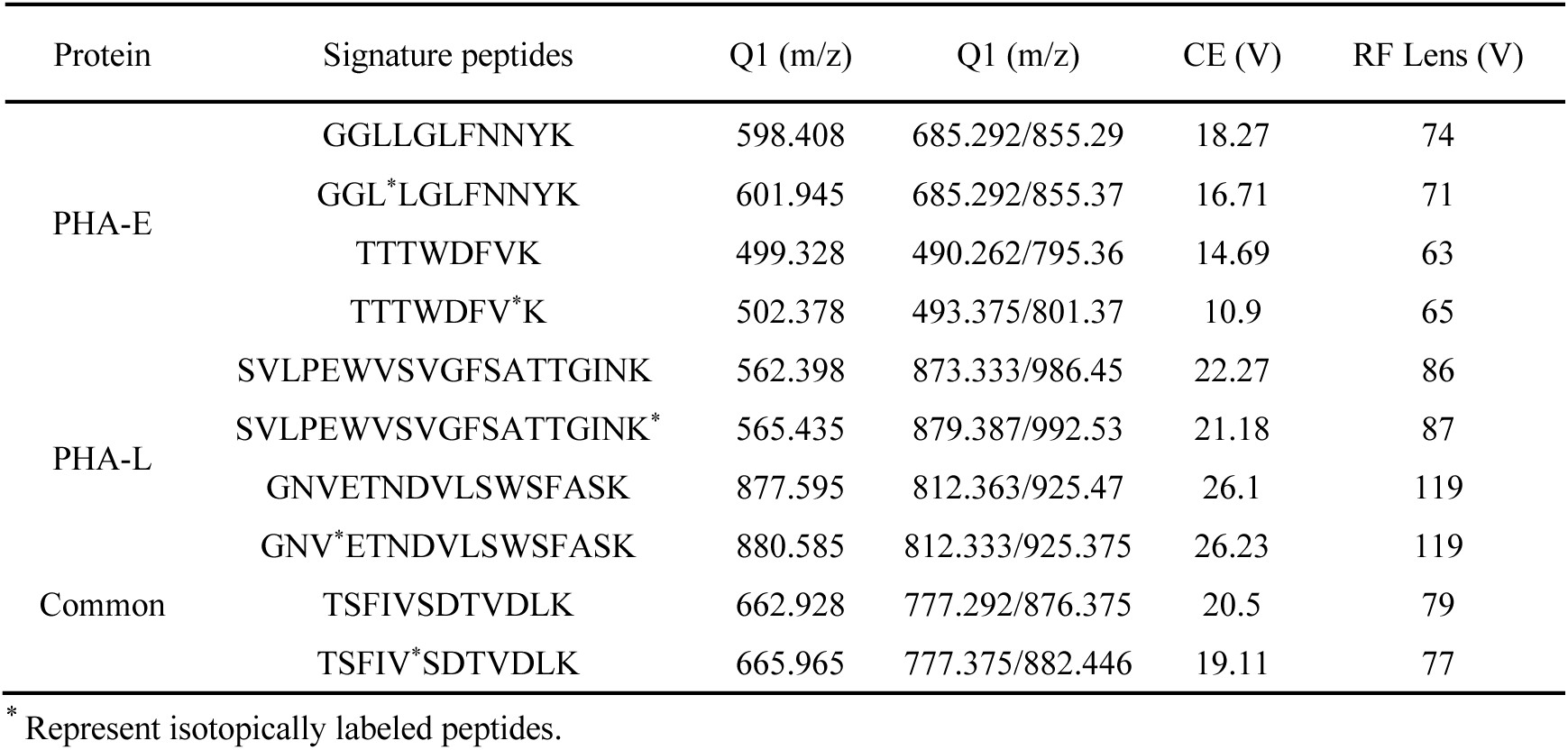
MRM parameters for analyzing the signature peptides.

### 3.2. Purity determination of the synthesized signature peptides using AAA-IDMS

To achieve absolute quantification of protein, AAA-IDMS is often used to quantify the exact mass fraction of the target peptide in the synthesized signature peptide. Val, Phe, Ile and Leu are commonly used amino acids in the AAA-IDMS method. In this study, these four amino acids were selected for quantitative analysis of the signature peptides. The key measurement parameters, including main condition of ionization, ion collision energy, and radio frequency (RF) voltage were optimized using the MRM mode of the MS and are shown in Table 2. In order to obtain the accurate mass fraction value, the acid hydrolysis time was optimized. From comparing ten different acid hydrolysis equilibrium time (0, 12, 24, 36, 48, 60, 72, 84, 96, 108 h), an optimized equilibrium time for each of the four signature peptides were determined. For example, the relative area ratios of Leu to ^13^C_5_-Leu, Phe to ^13^C_5_-Phe, Ile to ^13^C_5_-Ile, Val to ^13^C_5_-Val all increased quickly following the onset of hydrolysis, and they reached their peaks at 60h post hydrolysis for HIGIDVNSIR and 84h for GNVETNDVLSWSFASK, respectively (Fig. 2).

**Fig. 2.**
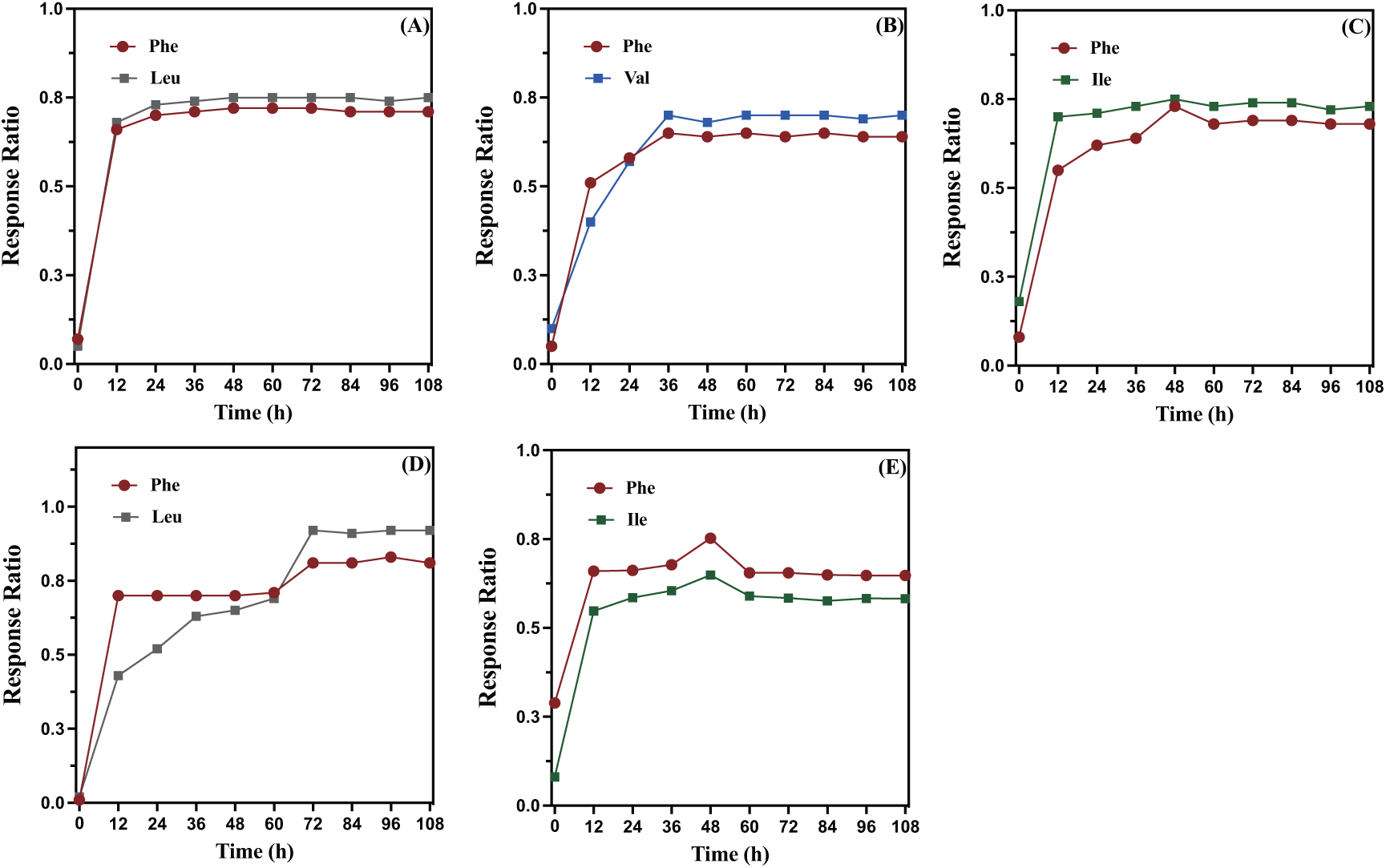
Optimization for the acid hydrolysis time. (A) GGLLGLFNNYK; (B)TTTWDFVK;(C) TSFIVSDTVDLK; (D) GNVETNDVLSWSFASK;(E) SVLPEWVSVGFSATTGINK.

**Table 2.** The MRM parameters of AAA-IDMS analysis.

| Compound | Precursor (m/z) | Product (m/z) | Collision Energy(V) | RF Lens(V) |
| --- | --- | --- | --- | --- |
| Ile | 131.962 | 86.196 | 10.23 | 31 |
| Ile* | 137.962 | 91.208 | 10.23 | 32 |
| Leu | 131.967 | 85.967 | 9.45 | 32 |
| Leu* | 138.05 | 91.05 | 9.8 | 32 |
| Phe | 165.962 | 120.196 | 14.51 | 38 |
| Phe* | 175.05 | 128.125 | 15.72 | 39 |
| Val | 117.967 | 71.967 | 10.21 | 30 |
| Val* | 123.017 | 75.967 | 14.76 | 30 |

\* Represent isotopically labeled AA.

Next, the isotope dilution internal standard method based on amino acid analysis was used for quantitative analysis. Various mass ratios (0.5:1, 0.75:1, 1:1, 1.25:1, 1.5:1) for 4 standard amino acids to labeled amino acids were tested. The actual mass ratio of standard amino acids to labeled amino acids served as the horizontal coordinate. The standard curve was constructed using the peak area ratio between the standard amino acid and the characteristic ion of the labeled amino acid as the ordinate. The purity of the synthesized peptides was then calculated based on this standard curve, and the results are presented in Table 3. It is noteworthy that the purity of the synthesized signature peptides ranged from 70% to 86%, indicating a considerable degree of variance. The conventional LC-MS/MS method overlooks the impurity of these signature peptides during measurement, making it susceptible to bias in comparison to our method.

**Table 3.**
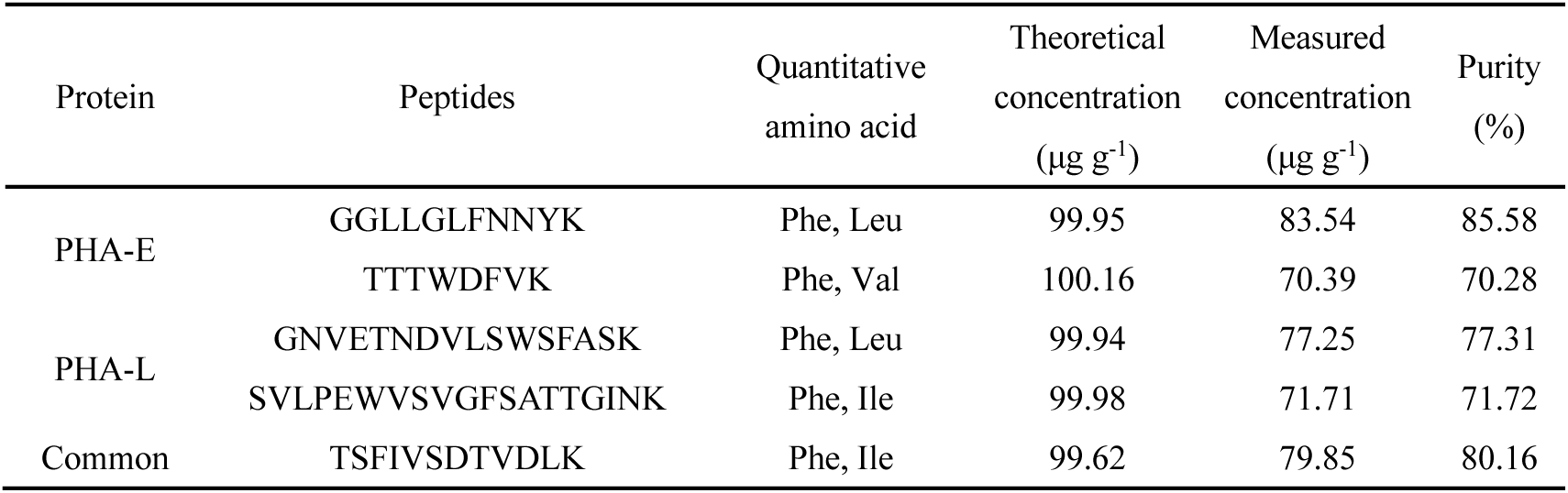
Purity of synthesized signature peptides as determined by AAA-IDMS (n=3).

### 3.3. Determination of an optimal total protein extraction protocol

For real biological samples, the selection of an appropriate total protein extraction protocol is critical for accurate quantification of specific proteins. In this study, we compared three different methods for extracting total protein from 50 mg of ground bean seeds, which included TCA-acetone precipitation method, TCA-acetone-phenol method, and phenol extraction methods. As shown in Fig. S2, the BCA assay revealed that the TCA-acetone method yielded the highest protein content at 5.5 mg, while the phenol method resulted in the lowest protein content at 0.4 mg. Moreover, as illustrated in Fig. 3, only TCA-acetone can successfully detect these five peptides and show high signal values.

**Fig. 3.**
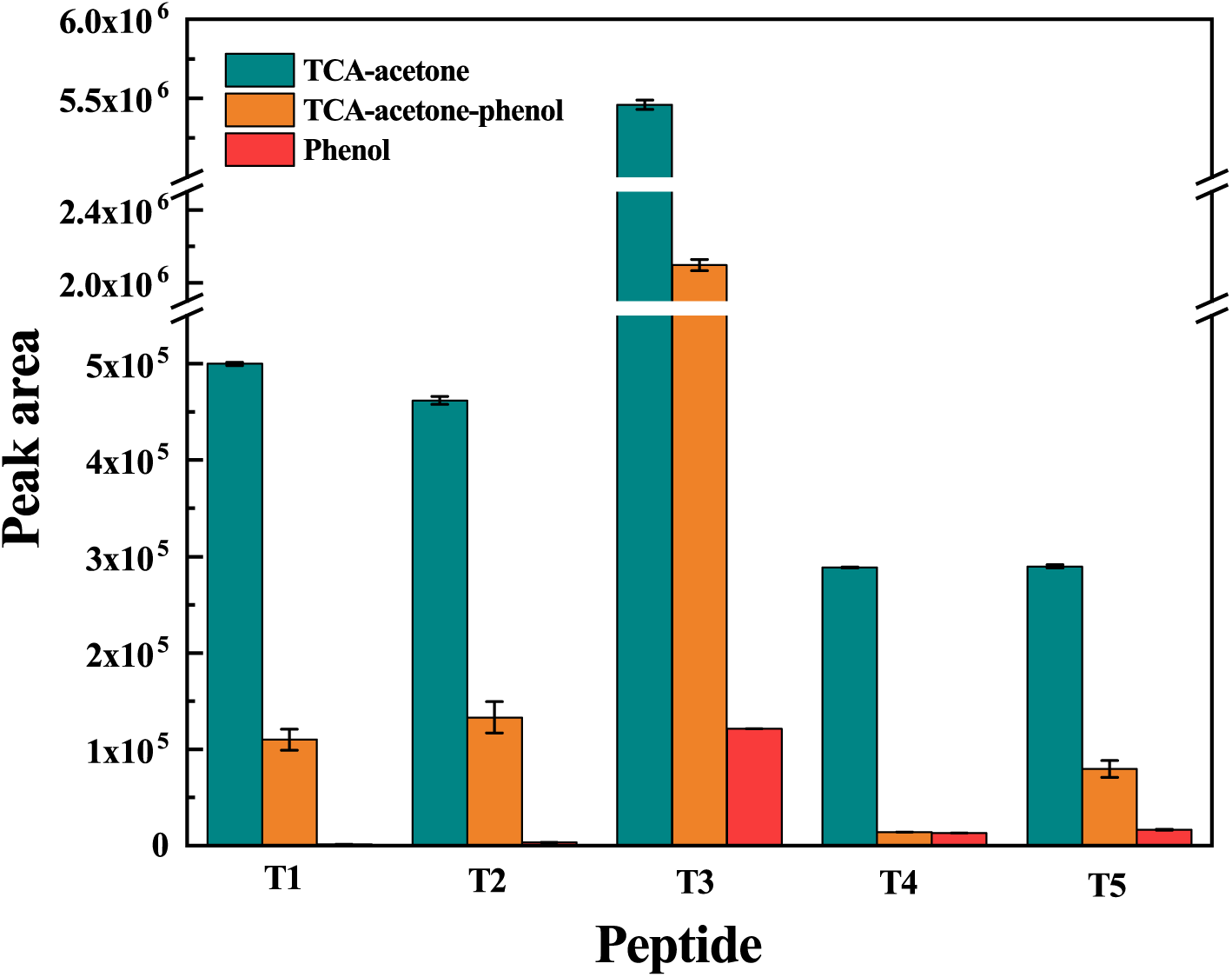
Comparison of mass spectra of three extraction methods (n=3). *T1 and T2 are PHA-E specific, T1: GGLLGLFNNYK T2: TTTWDFVK; T4 and T5 are PHA-L specific, T4: GNVETNDVLSWS, T5: SVLPEWVSVGFSATTGINK; T3: TSFIVSDTVDLK is common to both PHA-E and PHA-L.

### 3.4. Quantitative analysis for real samples

Utilizing the established methods, we conducted quantitative analysis of PHA in six distinct varieties of common beans. Standard curves were created for each of the five signature peptides, each consisting of six points, all demonstrating excellent linearity (R^2^ > 0.999, Fig. 4). As shown in Table 4, the quantification results for both PHA-E and PHA-L showed remarkable consistency when employing two different subunit-specific signature peptides. Furthermore, the total PHA content, quantified using the common signature peptide, closely matched the sum of the specific PHA-E and PHA-L contents. Across the six varieties, the total PHA content exhibited only medium level of variation, ranging from 41.69 ng mg^-1^ to 51.96 ng mg^-1^. However, the content of each PHA subunit displayed greater genotypic variance. JH-301 had the lowest PHA-E content but the highest PHA-L content. A consistent trend among the six varieties was the higher abundance of PHA-L compared to PHA-E. This result aligns with the gene expression data obtained from transcriptomic sequencing of the common bean seeds 20 days post anthesis, in which the Reads Per Kilobase Million (RPKM) values of *PHA-E* and *PHA-L* are 66,160 and 96,037, respectively (unpublished). These results thus affirm the unique resolution and accuracy of our method.

**Fig. 4.**
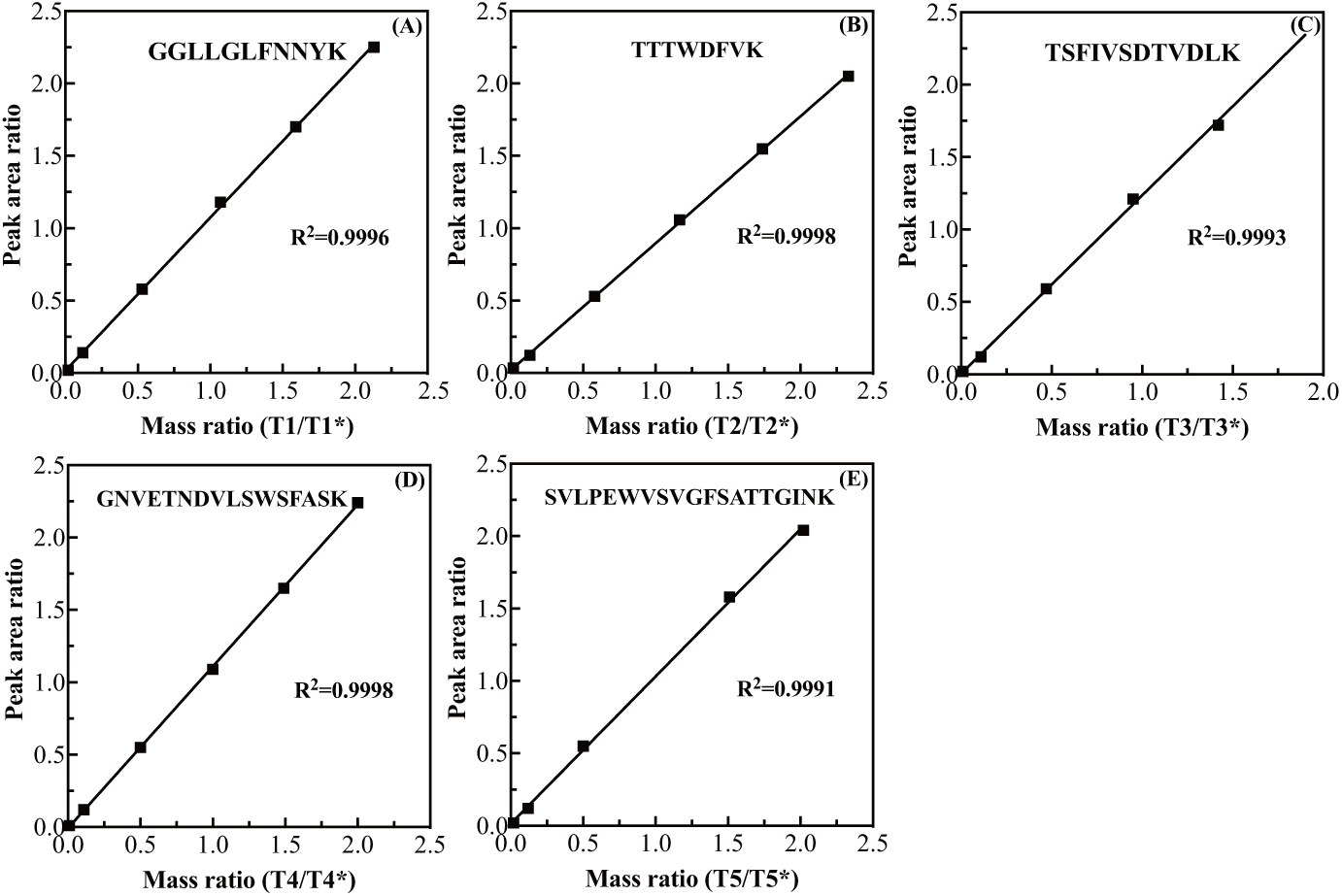
Standard curves of five signature peptides for PHA quantification. (A) T1: GGLLGLFNNYK; (B) T2: TTTWDFVK; (C) T3: TSFIVSDTVDLK; (D) T4: GNVETNDVLSWS; (E) T5: SVLPEWVSVGFSATTGINK. T1 and T2 are PHA-E specific, T4 and T5 are PHA-L specific, while T3 is common to both. *Represent isotopically labeled peptides.

**Table 4.** PHA quantification results for six common bean varieties (n=3).

| Variety | PHA-E (ng mg<sup>-1</sup>) |  | PHA-L (ng mg<sup>-1</sup>) |  | PHA-E+ L (ng mg<sup>-1</sup>) |
| --- | --- | --- | --- | --- | --- |
|  | T1 | T2 | T4 | T5 | T3 |
| JH-40 | 11.16 | 11.29 | 30.31 | 30.21 | 42.06 |
| JH-120 | 17.82 | 19.81 | 25.78 | 25.58 | 44.76 |
| JH-266 | 17.32 | 15.64 | 28.6 | 26.47 | 47.20 |
| JH-227 | 14.37 | 15.52 | 24.11 | 23.23 | 41.69 |
| JH-231 | 15.33 | 15.74 | 27.70 | 26.25 | 45.23 |
| JH-301 | 8.05 | 8.09 | 45.00 | 42.70 | 51.96 |

\*T1 and T2 are PHA-E specific, T1: GLLGLFNYYK T2: TTTWDFVK; T4 and T5 are PHA-L specific, T4: GNVETNDVLSWS, T5: SVLPEWVSVGFSATTGINK; T3: TSFIVSDTVDLK is common to both PHA-E and PHA-L.

### 3.5. Method Validation

Finally, the recovery, LOD, LOQ, intra-day precision, and inter-day precision were determined to validate the sensitivity and reliability of the developed method (Table 5). When the signal-to-noise ratios were set at 3 and 10, the detection limits for the five signature peptides ranged from 2.32 ng/mg to 12.25 ng/mg, while the quantitation limit ranged from 7.73 ng/mg to 40.82 ng/mg. Using the cultivar JH-40 as the material, the intra-day precision, represented as RSD values, varied between 0.23% (T2) to 3.45% (T5) for the subunit-specific signature peptides, and was low (0.63%) for the peptide common to both subunits (T3). The relatively high RSD observed for T5 may be attributed to its longer length, making it more susceptible to unfavorable analytical behaviors, such as hydrophobicity-related peptide adsorption or peak tailing during reversed-phase liquid chromatography (Klont et al., 2019). A similar level of inter-day precision was observed for each of the five signature peptides. Collectively, these results affirm they high sensitivity and repeatability of our established method for PHA content quantification in biological samples.

**Table 5.** Results of method validation (n=3)

| Peptides* | LOD<br>(ng g<sup>-1</sup>) | LOQ<br>(ng g<sup>-1</sup>) | Precision (%) |  | Recovery (%) | Standard curve |
| --- | --- | --- | --- | --- | --- | --- |
|  |  |  | Intra-day | Inter-day |  |  |
| T1 | 10.46 | 34.86 | 1.09 | 1.31 | 97.48~101.7 | y = 1.056x+0.017 |
| T2 | 7.06 | 23.52 | 0.23 | 0.44 | 94.18~95.7 | y = 8.768x+0.0185 |
| T3 | 2.32 | 7.73 | 0.63 | 0.93 | 104.47~101.04 | y = 1.0663x+0.0014 |
| T4 | 8.12 | 27.06 | 1.38 | 1.07 | 99.02~101.26 | y = 1.1197x-0.0096 |
| T5 | 12.25 | 40.82 | 3.45 | 3.27 | 94.94~100.08 | y = 1.0933x-0.0017 |

## 4 Conclusion

A specific and sensitive LC-MS/MS method has been developed, serving as a robust tool for precise quantitative analysis of PHA, a major antinutritional factor and allergenic protein in common beans. By identifying subunit-specific signature peptides, this method, to the best of our knowledge, enables the first-time differentiation of the highly similar PHA-E and PHA-L chains. In comparison to hemagglutination and ELISA assays, the use of specific signature peptides minimizes the risk of cross-reactivity, reducing the likelihood of false positives. The new method also overcomes the current scarcity of commercially available ELISA kit dedicated to PHA detection. The exemplary performance of this method is evident through its sensitive detection (2.32 ng mg^-1^) and quantitation (7.73 ng mg^-1^) limits, substantial recovery (94.18-104.47%), and excellent repeatability (RSD < 3.45%, in most cases < 1.5%). Furthermore, a comparative study highlights the TCA-acetone-based method for protein extraction as the recommended choice for sample pre-treatment. Effectively employed for the quantification of PHA in six common bean varieties, the established LC-MS/MS method not only highlights genotypic variations in PHA content but also discerns a consistent trend of greater abundance in PHA-L over PHA-E proteins in late-stage (20 days post-anthesis) developing seeds. This method stands to significantly enhance the identification and measurement of PHAs in the agriculture and food industry.

## CRediT authorship contribution statement

**Lan Li:** Conceptualization, Funding acquisition, Formal analysis, Writing – original draft, Writing – review & editing. Methodology. **Zhanying Chu:** Methodology, Validation. **Kang Ning:** Conceptualization. **Manman Zhu**: Methodology. **Rui Zhai**: Conceptualization, Formal analysis, Project administration, Supervision, Writing – review & editing. **Pei Xu:** Writing – review & editing, Writing – original draft, Visualization, Supervision, Resources, Project administration, Methodology, Conceptualization.

## Data availability

Data will be made available on request.

