## supplement figure and table for "A precise IDMS-based method for absolute quantification of phytohemagglutinin, a major antinutritional component in common bean"

**Table S1.** Thirteen candidate signature peptides and their characteristics obtained through bioinformatic analysis. The data of the five best-performing peptides are shown in bold and italic.

| Protein | Peptides | No. | Amino acid number | $MH^{+1}$ | $MH^{+2}$ | $MH^{+3}$ |
|---|---|---|---|---|---|---|
| PHA-E | LTNVNDNGEPTLSSLGR | T6 | 17 | 1786.893 | 893.9465 | 596.2977 |
| | ***GGLLGLFNNYK*** | ***T1*** | ***11*** | ***1195.647*** | ***598.3235*** | ***399.2157*** |
| | ***TTTWDFVK*** | ***T2*** | ***8*** | ***997.4989*** | ***499.2495*** | ***333.1663*** |
| | GENAEVLITYDSSTK | T7 | 15 | 1626.7857 | 813.8929 | 542.9286 |
| | SVLPEWVIVGFTATTGITK | T8 | 19 | 2019.1161 | 1010.0581 | 673.7054 |
| | LTNLNGNGEPR | T9 | 11 | 1184.6018 | 592.8009 | 395.5339 |
| PHA-L | DWDPTER | T10 | 7 | 918.3952 | 459.6976 | 306.7984 |
| | HIGIDVNSIR | T11 | 10 | 1123.6218 | 562.3109 | 375.2073 |
| | ***SVLPEWVSVGFSATTGINK*** | ***T5*** | ***19*** | ***1992.0437*** | ***996.5219*** | ***664.6812*** |
| | ***GNVETNDVLSWSFASK*** | ***T4*** | ***16*** | ***1753.8392*** | ***877.4196*** | ***585.2797*** |
| | LSDGTTSEGLNLANLVLNK | T12 | 19 | 1959.0393 | 980.0197 | 653.6798 |
| Common | ***TSFIVSDTVDLK*** | ***T3*** | ***12*** | ***1324.6995*** | ***662.8498*** | ***442.2332*** |
| | FNETNLILQR | T13 | 10 | 1247.6743 | 624.3372 | 416.5581 |

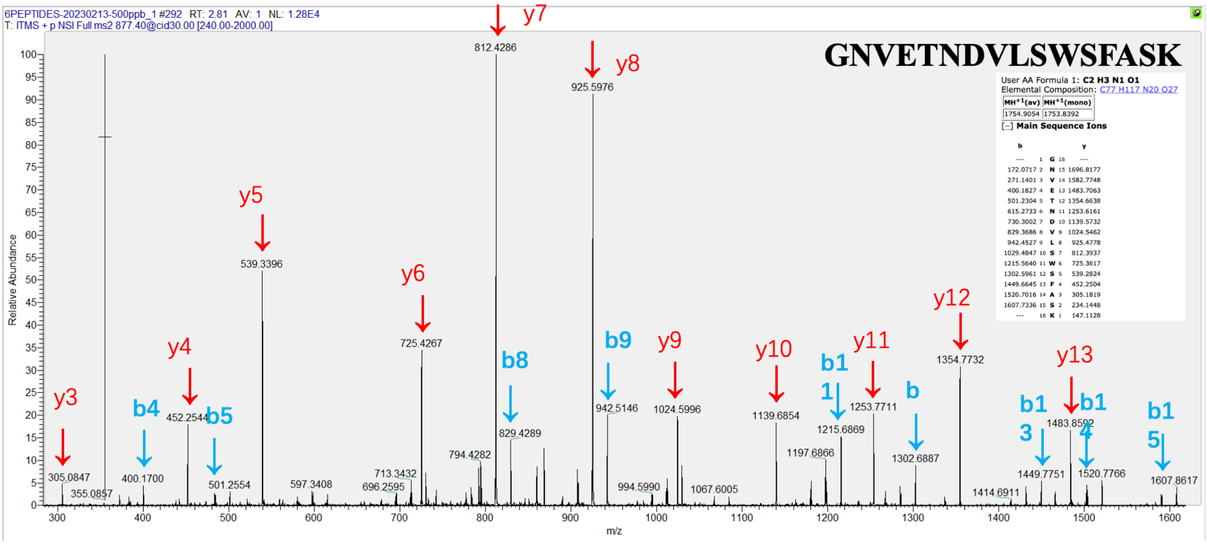

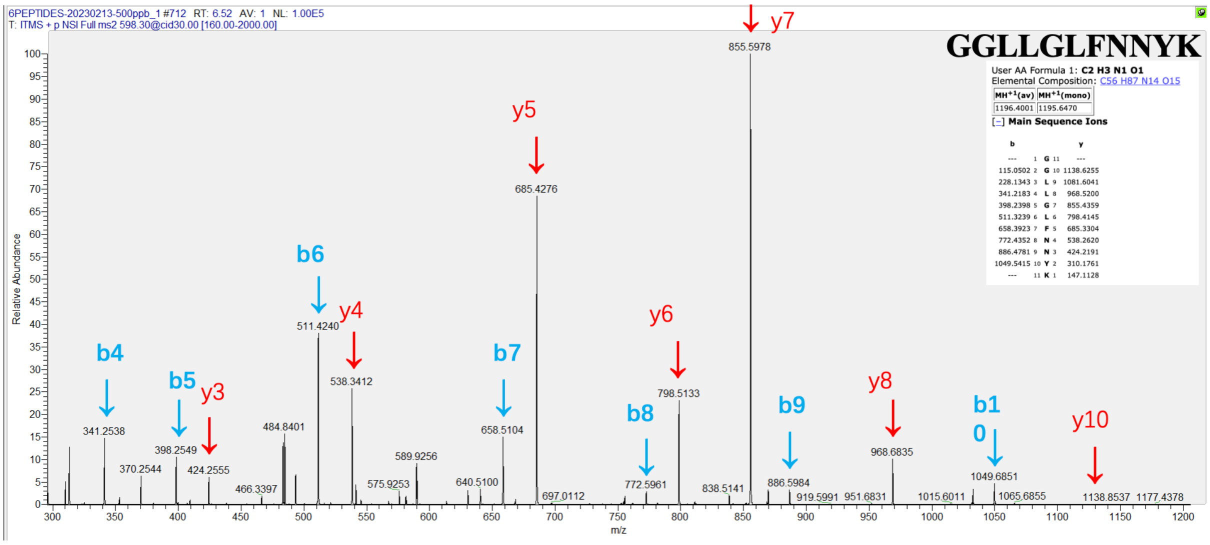
6PEPTIDES-20230213-500ppb_1 #712 RT: 6.52 AV: 1 NL: 1.00E5
T: ITMS + p NSI Full ms2 598.30@cid30.00 [160.00-2000.00]
GGLLGLFNNYK
User AA Formula 1: C2 H3 N1 O1
Elemental Composition: C56 H87 N14 O15
MH+1(av) MH+1(mono)
1196.4001 1195.6470
Main Sequence Ions
Relative Abundance
m/z
y7 y5 y4 y3 y6 y8 y10
b4 b5 b6 b7 b8 b9 b10
855.5978
685.4276
511.4240
538.3412
798.5133
341.2538
398.2549
424.2555
484.8401
658.5104
968.6835
1049.6851
1138.8537
1177.4378

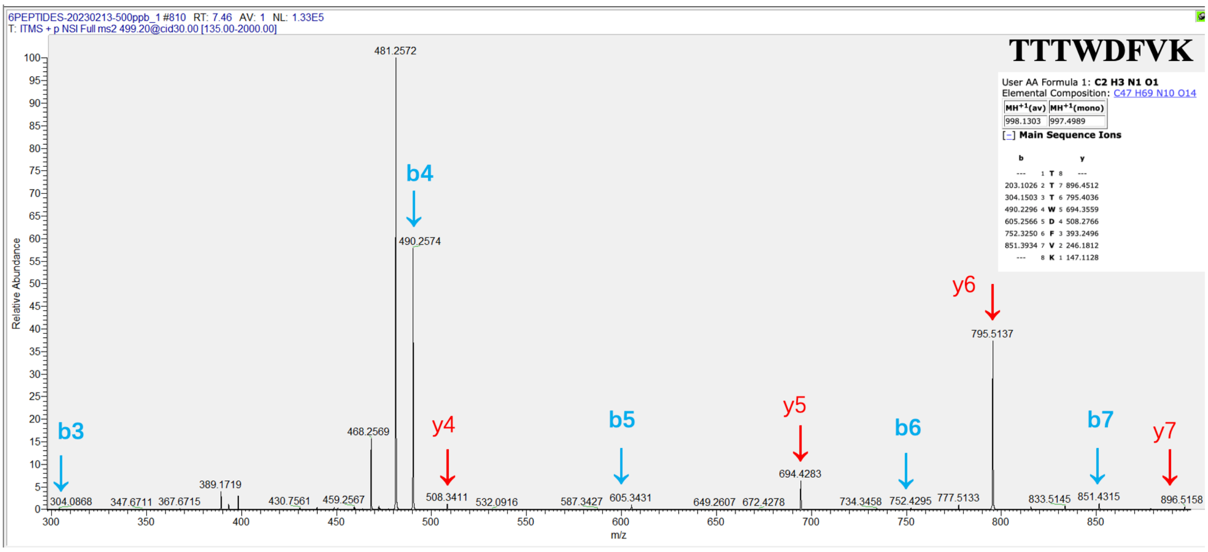
6PEPTIDES-20230213-500ppb_1 #810 RT: 7.46 AV: 1 NL: 1.33E5
T: ITMS + p NSI Full ms2 499.20@cid30.00 [135.00-2000.00]
TTTWDFVK
User AA Formula 1: C2 H3 N1 O1
Elemental Composition: C47 H69 N10 O14
MH+1(av) MH+1(mono)
998.1303 997.4989
Main Sequence Ions
Relative Abundance
m/z
b3 b4 b5 b6 b7
y4 y5 y6 y7
481.2572
490.2574
795.5137
468.2569
694.4283
304.0868
508.3411
605.3431
752.4295
851.4315
896.5158

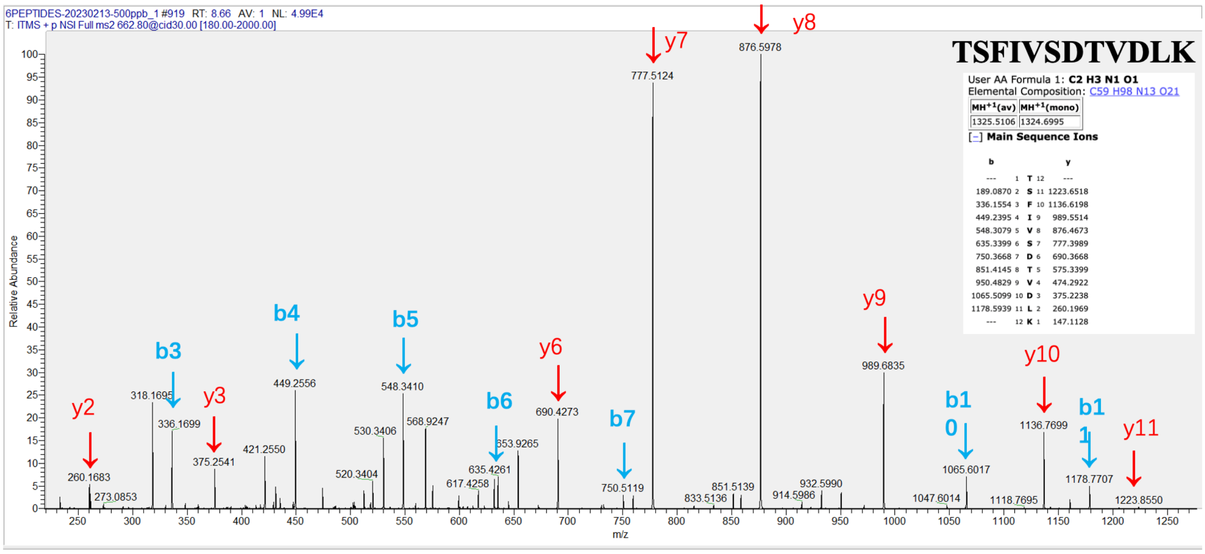
6PEPTIDES-20230213-500ppb_1 #919 RT: 8.66 AV: 1 NL: 4.99E4
T: ITMS + p NSI Full ms2 662.80@cid30.00 [180.00-2000.00]
TSFIVSDTVDLK
User AA Formula 1: C2 H3 N1 O1
Elemental Composition: C59 H98 N13 O21
MH+1(av) MH+1(mono)
1325.5106 1324.6995
Main Sequence Ions
Relative Abundance
m/z
y2 y3 y6 y7 y8 y9 y10 y11
b3 b4 b5 b6 b7 b10 b11
876.5978
777.5124
989.6835
449.2556
548.3410
690.4273
318.1695
336.1699
375.2541
260.1683
1136.7699
1178.7707
1065.6017
750.5119
1223.8550

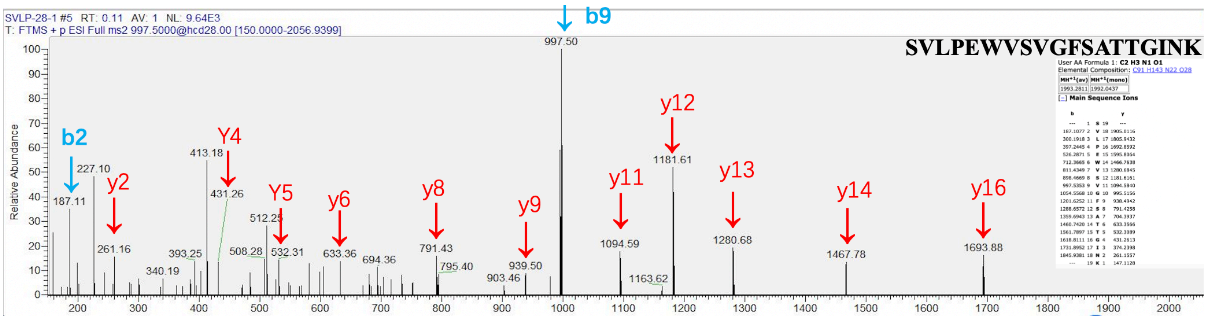


Fig. S1. Secondary structure diagrams of the signature peptides

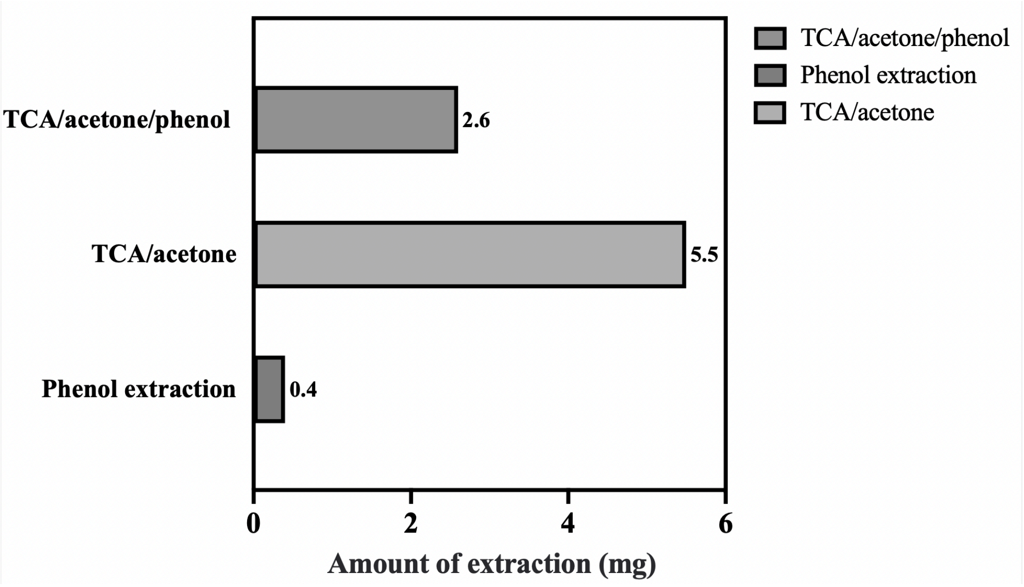


Fig. S2. Result of the BCA assay
